# Optogenetic investigation into the role of the subthalamic nucleus in motor control

**DOI:** 10.1101/2020.07.08.193359

**Authors:** Adriane Guillaumin, Gian Pietro Serra, François Georges, Åsa Wallén-Mackenzie

## Abstract

The subthalamic nucleus is important achieve intended movements. Loss of its normal function is strongly associated with several movement disorders. Classical basal ganglia models postulate that two parallel pathways, the direct and indirect pathways, exert opposing control over movement, with the subthalamic nucleus part of the indirect pathway through which competing motor programs are prevented. The subthalamic nucleus is regulated by both inhibitory and excitatory projections but experimental evidence for its role in motor control has remained sparse. The objective here was to tease out the selective impact of the subthalamic nucleus on several motor parameters required to achieve intended movement, including locomotion, balance and motor coordination. Optogenetic excitation and inhibition using both bilateral and unilateral stimulations of the subthalamic nucleus were implemented in freely-moving mice. The results demonstrate that selective optogenetic inhibition of the subthalamic nucleus enhances locomotion while its excitation reduces locomotion. These findings lend experimental support to basal ganglia models in terms of locomotion. However, further analysis of subthalamic nucleus excitation revealed grooming and disturbed gait. Selective excitation also caused reduced motor coordination, independent of grooming, in advanced motor tasks. This study contributes experimental evidence for a regulatory role of the subthalamic nucleus in motor control.

**Highlights:**

- Bilateral optogenetic excitation of the subthalamic nucleus in freely-moving mice reduces forward locomotion while optogenetic inhibition leads to its increase.
- Unilateral optogenetic excitation and inhibition of the subthalamic nucleus cause opposite rotational behavior.
- Bilateral optogenetic excitation, but not inhibition, of the subthalamic nucleus induces jumping and self-grooming behavior.
- Engaged in advanced motor tasks, bilateral optogenetic excitation causes mice to lose motor coordination.
- The results provide experimental support for predictions by the basal ganglia motor model on the role of the subthalamic nucleus in locomotion, and identifies a causal role for the subthalamic nucleus in self-grooming.

## Introduction

The subthalamic nucleus (STN) is a small, bilaterally positioned structure which exerts regulatory influence over voluntary movement. Consequently, damage or dysregulation of the STN is strongly associated with motor dysfunction and movement disorder. For example, unilateral damage to the STN causes strongly uncontrolled movements, so called hemiballismus (Hamada & DeLong, 1992), while degeneration of the STN is associated with supranuclear palsy and Huntington’s disease (Dickson et al., 2010; Lange et al., 1976). Further, hyperactivity of STN neurons is a pathological hallmark of Parkinson’s disease (PD) (Albin et al., 1989; DeLong, 1990). Surgical lesioning of the STN, so called subthalamotomy, improves motor symptoms in PD (Heywood & Gill, 1997; Parkin et al., 2001), and so does deep brain stimulation (DBS), an electrical method in which high-frequency stimulation electrodes, when positioned in the STN, can correct its aberrant activity (Benazzouz et al., 2000; Filali et al., 2004). DBS of the STN (STN-DBS) successfully alleviates motor symptoms in advanced-stage PD (Benabid, 2003; Benabid et al., 2009). The high success rate of these clinical interventions confirms the critical role of the STN, yet its regulatory role over different motor parameters required to achieve willed movement remains to be fully established.

The critical role of the STN in motor function is commonly explained by its connectivity on the neurocircuitry level in which it forms an integral part of the basal ganglia. According to the classical basal ganglia model, two pathways, the so called direct and indirect pathways, exert opposing control over motor behavior via the thalamus (Albin et al., 1989; Alexander & Crutcher, 1990; DeLong, 1990; Graybiel et al., 1994; Smith et al., 1998). Via the direct pathway (cortex - striatum - internal segment of the globus pallidus (GPi) – thalamus), intended movements are promoted, while via the indirect pathway (cortex - striatum - external segment of the globus pallidus (GPe) – STN – GPi - thalamus), competing motor programs are suppressed and stop signals mediated (Nambu et al., 2002; Sano et al., 2013; Schmidt & Berke, 2017). In addition to the indirect pathway, the STN is directly regulated by the cortex in the so called hyperdirect pathway. The STN itself is excitatory, using glutamate as neurotransmitter. Extensive STN projections reach the GPi and substantia nigra pars reticulata (SNr), and the STN is also reciprocally connected to the GPe. This organization effectively means that the STN activity is regulated by both excitatory (cortex) and inhibitory (GPe) projections. When STN neurons are activated, they contribute to the suppression and stopping of movement by counteracting the movement-promoting activities of the direct pathway (Féger et al., 1994; Gradinaru et al., 2009; Mouroux & Féger, 1993; Sanders & Jaeger, 2016; Sugimoto & Hattori, 1983)

This model of STN’s role in motor regulation via the basal ganglia pathways has been formed primarily based on anatomical projection patterns and functionally by lesion and high-frequency stimulation in experimental animals, as well as studies of human symptoms, primarily in PD (Alkemade et al., 2015; Benabid, 2003; Benabid et al., 2009; Benazzouz et al., 2000; Centonze et al., 2005; Hamada & DeLong, 1992; Hamani, 2004; Haynes & Haber, 2013; Heywood & Gill, 1997; Parent & Hazrati, n.d.; Parkin et al., 2001; Rizelio et al., 2010). However, experimental evidence has remained sparse, in particular in terms of STN excitation. In addition, studies based on human pathological conditions suffer from complexity in terms of confounding factors, such as multiple symptom domains and pharmacological treatments. Another complicating issue is the fact that data derived from DBS-studies can be difficult to fully understand since the underlying mechanisms of this method have remained unresolved (Chiken & Nambu, 2016; Florence et al., 2016; Vitek, 2002). Experimental animals in which carefully controlled conditions can be achieved and functional output reliably recorded are therefore essential to increase the understanding of the roles that distinct structures play in circuitry and behavioral regulation. Indeed, several studies in rodents have supported the hypothesis that removal of STN activity enhances movement. Both STN-lesions and STN-selective gene knockout of the *Vesicular glutamate transporter 2* (Vglut2), encoding the transporter that packages glutamate for synaptic release, lead to behavioral phenotypes of enhanced locomotion (Centonze et al., 2005; Rizelio et al., 2010; Schweizer et al., 2014, 2016). However, lesioning and gene-targeting methods suffer from drawbacks and do not allow the analysis of how the intact STN regulates movement. Further, gene-targeting during brain development might cause structural and functional adaptations that likely contribute to behavioral phenotypes in adulthood.

During the past decade, the use of optogenetics in freely-moving rodents has enabled more advanced possibilities for decoding the role of distinct neuronal structures and circuitries in behavioral regulation. By allowing spatially and temporally precise control over neural activity using excitatory and inhibitory opsins (Deisseroth, 2011), ground-breaking discoveries have been made, not least in terms of motor control. This includes the experimental validation of the classical model of the basal ganglia using excitatory and inhibitory opsins in striatal neurons, verifying the opposing roles of the direct and indirect pathways in regulation of movement (Kravitz et al., 2010). More recently, optogenetic dissection of neurons in the globus pallidus verified their role in motor control (Tian et al., 2018). In terms of optogenetics in the STN, the strong interest in understanding the mechanisms of STN-DBS has led to a focus on PD models. However, somewhat surprisingly, the first study addressing the STN using optogenetics did not find any effect on recovery in a PD model using either excitatory or inhibitory opsins in the STN itself, while identifying motor improvement only upon excitation of the cortico-subthalamic pathway (Gradinaru et al., 2009), a finding subsequently confirmed (Sanders & Jaeger, 2016). In contrast, a more recent study reported improved motor function in a rodent PD-model upon optogenetic inhibition of the STN structure (Yoon et al., 2014). Thus, contradicting results have been obtained with optogenetic excitation and inhibition of the STN structure. Several factors might contribute to the differences in findings, including the transgenic line implemented to drive the expression of the opsin constructs, the behavioral setup, the PD model, and more.

While the use of optogenetics in various rodent PD models have begun to answer fundamental questions around STN-DBS mechanisms, the natural role of the STN in regulating different aspects of motor behavior during non-pathological conditions has remained poorly explored. However, a basic understanding of STN’s regulatory role in movement should prove useful for advancing both pre-clinical and clinical knowledge. The objective of the present study was to tease out the impact of the STN on several motor parameters required to achieve intended movement, including locomotion, balance and motor coordination. To ensure opsin expression in the STN structure, Pitx2-Cre mice were used based on previous validations of the Pitx2-Cre transgene as a selective driver of floxed opsin constructs in the STN (Schweizer et al., 2014; Viereckel et al., 2018). Mice expressing excitatory or inhibitory opsins selectively in the STN were analysed both upon bilateral and unilateral photostimulations. The findings reported contribute to increased understanding of motor control by both confirming and challenging the role of the STN as predicted by the basal ganglia motor model.

## Material and methods

### Animal housing and ethical permits

Pitx2-Cre_C57BL/6NTac transgenic mice were bred in-house and housed at the animal facility of Uppsala University before and during behavioral experiments or at University of Bordeaux after virus injections for in vivo electrophysiological experiments. Mice had access to food and water ad libitum in standard humidity and temperature conditions and with a 12 hour dark/light cycle. PCR analyses were run to confirm the genotype of Pitx2-Cre positive mice from ear biopsies. All animal experimental procedures followed Swedish (Animal Welfare Act SFS 1998:56) and European Union Legislation (Convention ETS 123 and Directive 2010/63/EU) and were approved by local Uppsala or Bordeaux (N°50120205-A) Ethical Committee.

### Stereotaxic virus injection and optic cannula implantation

#### Virus injection

Stereotaxic injections were performed on anesthetized Pitx2-Cre mice maintained at 1.4-1.8 % (0.5-2 L/min, isoflurane-air mix v/v). Before starting the surgery and 24h post-surgery mice received subcutaneous injection of Carprofen (5 mg/Kg, Norocarp). A topical analgesic, Marcain (1.5 mg/kg; AstraZeneca) was locally injected on the site of the incision. After exposing the skull, holes were drilled in the skull for virus injections and skull screws implantations for behavioral experiments. Mice were injected in the STN bilaterally with a virus containing either a Cre-dependent channelrhodopsin (rAAV2/EF1a-DIO-hChR2(H134R)-eYFP) (mice referred to as Pitx2/ChR2 mice), a Cre-dependent archaerhodopsin (rAAV2/EF1a-DIO-eArch3.0-eYFP) (mice referred to as Pitx2/Arch) or an empty virus carrying only a fluorescent protein (rAAV2/EF1a-DIO-eYFP) (mice referred to as Pitx2/eYFP-C for controls within the ChR2 group, and Pitx2/eYFP-A for controls within the Arch group): x 3.8×1012 virus molecules/mL, 2.7×1012 virus molecules/mL, 4.6×1012 virus molecules/mL, respectively (UNC Vector Core, Chapel Hill, NC, USA). The following coordinates were used (adapted from Paxinos and Franklin, 2013): anteroposterior (AP) = −1.90 mm, mediolateral (ML) = +/− 1.70 mm from the midline. 250 nL of virus was injected with a NanoFil syringe (World Precision Instruments, Sarasota, FL, USA) at two dorsoventral levels (DV) = −4.65 mm and −4.25 mm from the dura matter at 100 nL.min-1.

#### Optic cannula implantation

Optic cannulas (MFC_200/245-0.37_5mm_ZF1.25_FLT, Doric Lenses) were implanted directly after the virus injections. Two skull screws were implanted in the skull to hold the optic cannula-cement-skull complex. Primers (OptibondTM FL, Kerr) were then applied and harden with UV light. Optic cannulas were implanted bilaterally above the STN and fixed with dental cement, coordinates: AP = −1.90 mm, ML = +/− 1.70 mm from the midline DV = −4.30 mm. 1 mL of saline was injected subcutaneously at the end of the surgery.

### Single-cell extracellular recordings

#### Surgery

In vivo single cell extracellular recordings started at least 4 weeks after virus injections. Mice were anesthetized with a mix isoflurane-air (0.8-1.8 % v/v) and placed in a stereotaxic apparatus. Optic fiber, optrode and glass micropipettes coordinates were AP = −1.90 mm, ML = +/− 1.70 mm and DV = −4.30 mm for the STN and respectively anterior and central GP: AP = −0.11 mm relative to bregma, ML = +/− 1.50 mm from sagittal vein, DV = from −3.00 to −5.50 mm depth and AP = −0.71 mm relative to bregma, ML = +/− 1.80 mm, DV:−3.00/−4.20 mm.

#### STN optotagging

A custom-made optrode was made with an optic fiber (100 µm diameter, Thorlabs) connected to a laser (MBL-III-473 nm-100 mW laser, CNI Lasers, Changchun, China) mounted and glued on the recording glass micropipette which was filled with 2% pontamine sky blue in 0.5□µM sodium acetate (tip diameter 1-2 µm, resistance 10-15 MΩ). The distance between the end of the optic fiber and the tip of the recording pipette varied between 650 nm and 950 nm. Extracellular action potentials were recorded and amplified with an Axoclamp-2B and filtered (300□Hz/0.5□kHz). Single extracellular spikes were collected online (CED 1401, SPIKE2; Cambridge Electronic Design). The laser power was measured before starting each experiment using a power meter (Thorlabs). The baseline was recorded for 100 seconds for each neuron before starting any light stimulation protocols which were set and triggered with Spike2 software. Light protocol consisted in a peristimulus time histogram (PSTH, 0.5 Hz, 5 ms bin width, 5-8 mW) for at least 100 seconds.

#### GP recordings

An optic fiber was placed above the STN and GP neurons were recorded. For each GP neurons, a PSTH was recorded upon STN optogenetic stimulation (PSTH 0.5 Hz, 5 ms pulses, 5-8 mW) for at least 100 seconds. Once the neuronal activity returned to baseline the same light stimulation protocol used in behavioral experiments was applied for 100 seconds (20 Hz, 5 ms pulses, 5-8 mW, referred as “Behavioral protocol”). Neurons are considered as excited during the PSTH protocol when, following the light pulses centered on 0, the number of spikes/5ms bin is higher than the baseline (−500 ms to 0 ms) plus two times the standard deviation.

#### Histology

At the end of each experiment, a deposit was made at the last recording coordinate by pontamine sky blue electrophoresis (−20□μA, 25□min). Brains were then collected after 0.9% NaCl and 4% PFA transcardial perfusion. 60 µm brain sections were cut with a vibratome at the levels of the STN and the GP, mounted with Vectashield medium (Vector Laboratories), coverslipped and visualized with an epifluorescent microscope to confirm eYFP expression, recording pipette and optic fiber positions.

### c-Fos activation

Pitx2/ChR2 and respective Pitx2/eYFP-C control mice were connected to the optic fiber and put in a neutral cage for 20 min before receiving 2 min light stimulation, 5 mW, 5 ms pulse duration, 20 Hz. Mice were perfused 90 min after the light stimulation.

### Histological analysis

Following behavioral tests, mice were deeply anesthetized and perfused transcardially with Phosphate-Buffer-Saline (PBS) followed by ice-cold 4% formaldehyde. Brains were extracted and 60 μm sections were cut with a vibratome. Immunohistochemistry for eYFP expression was run on sections from every mice and immunohistochemistry for c-Fos expression was run on sections of mice which underwent a c-Fos activation protocol. After mounting, slices were scanned with NanoZoomer 2-0-HT.0 (Hamamatsu) scanner and visualized with NDPView2 software (Hamamatsu).

Fluorescent immunohistochemistry was performed to reveal c-Fos expression and enhance the eYFP signal. After rinsing in PBS, sections were incubated for 90 min in PBS 0.3% X-100 Triton containing 10% blocking solution (normal donkey serum) followed by an incubation with primary antibodies, diluted in 1% normal donkey serum in PBS, overnight at 4°C (chicken anti-GFP 1:1000, cat. no. ab13970, Abcam; rabbit anti-c-Fos 1:800, cat. no. 226003, Synaptic System). The next day, sections were rinsed in PBS plus 0.1% Tween-20 solution and incubated for 90 min with secondary antibodies diluted in PBS (Cy3 donkey anti-rabbit 1:1000; A488 donkey anti-chicken 1:1000, cat. no. 703-545-155, Jackson Immunoresearch). After rinsing in PBS plus 0.1% Tween-20 solution, sections were incubated for 30 min with DAPI diluted in distilled water (1:5000). Sections were mounted with Fluoromount Aqueous mounting medium (Sigma, USA) and cover-slipped.

### Behavioral testing

After approximately 4 weeks of recovering from surgery, mice were analyzed in validated behavioral paradigms, during which the same stimulation protocols were used: for Pitx2/ChR2 and respective controls, 473 nm light, 5 mW, 20 Hz, 5 ms pulse delivered by a MBL-III-473 nm-100 mW laser (CNI Lasers, Changchun, China); for Pitx2/Arch and respective controls, 532 nm continuous light, 10 mW, delivered by a MBL-III-532 nm-100 mW laser (CNI Lasers, Changchun, China). Duration and condition of stimulation were specified for each test. After completed behavioral tests, mice were sacrificed and brains analyzed histologically and for placement of optic cannulas. Mice in which the optic cannulas were not in correct position were excluded from the analysis.

### Habituation

Three weeks after surgery and before the first behavioral test, all mice were handled and habituated to the experimental room and to the optic cables to reduce the stress during the day of the experiment. Before each behavioral test, mice were acclimatized for 30 minutes in the experimental room.

### Open field test

Mice were individually placed in neutral cages for three minutes in order to recover after connecting the optic cables. Mice were subsequently placed in the central zone of the open field arena and allowed to freely explore it for 5 min before starting the test. The open-field chamber consisted in a 50 cm, squared, transparent, plastic arena with a white floor that has been divided into a central zone (center, 25% of the total area) and a peripheral zone (borders). The open field test consisted of a 20 minutes session divided in four alternating 5-minutes trials (OFF-ON-OFF-ON). The patch cable was connected to a rotary joint, which was attached on the other end to a laser that was controlled by an Arduino Uno card. During the light ON trials, blue light was delivered according to the behavioral stimulation protocol. Total distance moved, speed, time spent and frequency in crossing to the center and body elongation, were automatically documented. Rearing, grooming and escape behaviors were manually recorded by an experimenter blind to the experimental groups using the EthoVision XT 12.0/13.0 tracking software (Noldus Information Technology, The Netherlands).

### Unilateral STN-stimulation in the open field

The test was performed in the same testing arena used for the open field test and consisted of two 10-minute phases, each divided in two 5-minute trials (OFF-ON). Light was delivered during ON trials according to the protocol. On “Phase 1” the photostimulation was randomly paired to one hemisphere while on “Phase 2” it was delivered to the contralateral hemisphere. Before “Phase 1”, mice were allowed to recover from handling in a neutral cage for 5 minutes, then they were placed in the center of the arena and allowed to explore it for 5 minutes before starting the test. Between the two phases the optic fiber was switched from one hemisphere to the contralateral one. Again mice were allowed to recover from handling in a neutral cage for 5 minutes and then placed in the center of the arena for “Phase 2”. Total distance moved, body rotations, time spent and frequency in crossing to the center of the arena were automatically recorded while rearing activity, grooming and escape behavior were manually scored by using the EthoVision XT 13.0 tracking software (Noldus Information Technology, The Netherlands).

### Rotarod

A fixed-speed rotarod assay was performed to investigate the ability of mice to maintain balance on a rotating rod (BIOSEB, Vitrolles, France). Before testing, all animals were trained for 4 days (“Training Days”) on different trials with fixed-speed protocols (6, 8, 12, 16 rpm). Each trial consisted of a maximum of 3 attempts. Mice were individually placed in a neutral cage to recover from connection for 15 min before starting the trial. Each trial was considered to have started after 5 s that the mouse was placed on the rotating drum and ended either when the mouse fell from the rod or after a total time of 120 s had elapsed. Resting time between trials was 5 min. Once an animal succeeded to stay on the rod for 120 s it was approved to the next trial according to the daily schedule. The “Pre Test” day consisted of 4 trials at a fixed speed of 16 rpm, with no light applied and served to test mice’s balance skills in absence of optogenetic stimulation. Only mice able to pass the “Pre Test” were tested on the next day. The “Test Day” was performed throughout 4 trials at a fixed speed of 16 rpm, with OFF trials (no photostimulation) alternated with ON trials (photostimulation). The laser was started 5 s after the mouse was placed on the rod, in conjunction with the start of each ON trial, and turned off when the animal fell or when the 120s time slice was over. Latency to fall was automatically scored by the Ethovision XT13.0 tracking software (Noldus Information Technology, The Netherlands). When more than one attempt was needed, or when an animal was not able to stay on the rod for 120 s, only the best performance among the 3 attempts was taken for statistical analysis.

### Beam walk test

Mice were trained to walk from a start position along a wooden round beam (80 cm of total length, 25, 21 and 15 mm diameter) to a goal cage. The beams were placed horizontally, elevated 40 cm from the bench, with one end held by a metal holder and the other one leaned against a neutral cage. The whole beam was divided in 3 parts: a “Starting Area” (SA, 10 cm length) which was brightly lit with a 60 W lamp to induce an aversive stimulus to move forward on the beam, a “Recording Area” (RA, 60 cm segment) where all the parameters were assessed, and a “Goal Zone” (GZ, 10 cm) attached to a cage (clean bedding and home cage nesting material) into which the mouse could escape. On the “Training Days” (TD) trials were performed on the 25, 21 (TD 1) and 15 mm (TD 2) diameter beams until two consecutive trials were performed with no stopping. Mice were individually placed in the goal cage for 15 min before starting the training in order to recover from optic cable connection and habituate to the goal cage. Subsequently mice were individually positioned on the SA and allowed to walk to the GZ. On the “Test” day, four different conditions were tested on a 15 mm diameter beam. Latency to cross the RA and number of hind foot slips were recorded and scored according to the pre-defined rating scale (Table after 1 and adapted for mice). The software EthoVision XT 13 was used to record the time to cross the RA, while the number of hind foot slips were recorded manually. Each trial session was started by the mouse entering the RA and ended by the mouse reaching the GZ or falling from the beam. Resting time in the goal cage lasted for 3 min between sessions. The four tested condition were: “Baseline” (BS), to measure mice balance skills in absence of photostimulation; “Alternated Light” (ALT), with photostimulation applied in alternated 15 cm-segments within the RA (OFF and ON sectors); “Continuous Light” (CONT), with the photostimulation applied across the whole RA, turned on by the mouse entering the RA and turned off by reaching the GZ or falling from the beam; “Prior Stimulation” (PRIOR), with the photostimulation delivered from 15 s before positioning the mouse on the SA of the beam until the GZ was reached or the mouse fell. For each condition two trials in which the mouse did not stall on the beam were averaged. The pre-defined rating scale “neuroscore” was designed by having the maximum score set to 10 points for each condition (1-5 points for latency to cross + 1-5 points for hind-foot slips). The lower the score, the better was the motor performance. See Supplementary Table 1 for scoring scale.

### Statistics

Results from all statistical analyses are shown in Supplementary Table 1. Data from behavioral test are expressed on the plots as means ± SEM and when necessary were averaged for the two ON and OFF trials. For statistical analysis Two tailed Wilcoxon matched-pairs test or RM Two-way ANOVA were performed followed by Bonferroni’s or Tukey’s multiple comparisons where appropriate.

Normal distribution of residuals was checked by using QQ plot and Shapiro-Wilk W test. Rank based z score transformation was applied prior the Two-way ANOVA analysis when more than two outliers were detected. For single cell extracellular recordings Friedman test was performed followed by Dunn’s multiple comparisons. For rotarod test and beam walk test, Friedman test was performed followed by Dunn’s multiple comparisons (GraphPad Prism version 7.00 for Windows, GraphPad Software, La Jolla California USA).

## Results

### Expression of optogenetic constructs in the STN of Pitx2-Cre mice with c-Fos induced upon photoexcitation confirms experimental setup

The *Pitx2* gene encodes a transcription factor essential for STN development and function (Martin et al., 2004; Schweizer et al., 2016; Skidmore et al., 2008). We previously verified that expression of the optogenetic ion channel Channelrhodopsin (ChR2) in the STN of Pitx2-Cre mice causes post-synaptic currents and glutamate release in STN target areas upon photostimulation (Schweizer et al., 2014; Viereckel et al., 2018), thus validating the use of optogenetics in Pitx2-Cre mice for the study of STN.

Here, to allow excitatory and inhibitory control of STN neurons, Pitx2-Cre mice were bilaterally injected with an adeno-associated virus (AAV2) containing a DNA-construct encoding either the excitatory ion channel hChR2 (Pitx2/ChR2 mice) or the inhibitory proton pump Archaerhodopsin eArch3.0 (Pitx2/Arch mice), both opsin constructs fused with the enhanced yellow fluorescent protein, eYFP (Figure 1A). As controls, separate cohorts of Pitx2-Cre mice were injected with AAV2 expressing the fluorescent reporter eYFP alone (same control virus but different cohorts of mice, thus referred to as Pitx2/eYFP-C mice for the controls to Pitx2/ChR2 mice; Pitx2/eYFP-A for the controls to Pitx2/Arch mice). In all mice, optical cannulas were placed above the STN allowing for photostimulation at 473 nm for Pitx2/ChR2 mice and 532 nm for Pitx2/Arch mice (Figure 1A).

**Figure 1.**
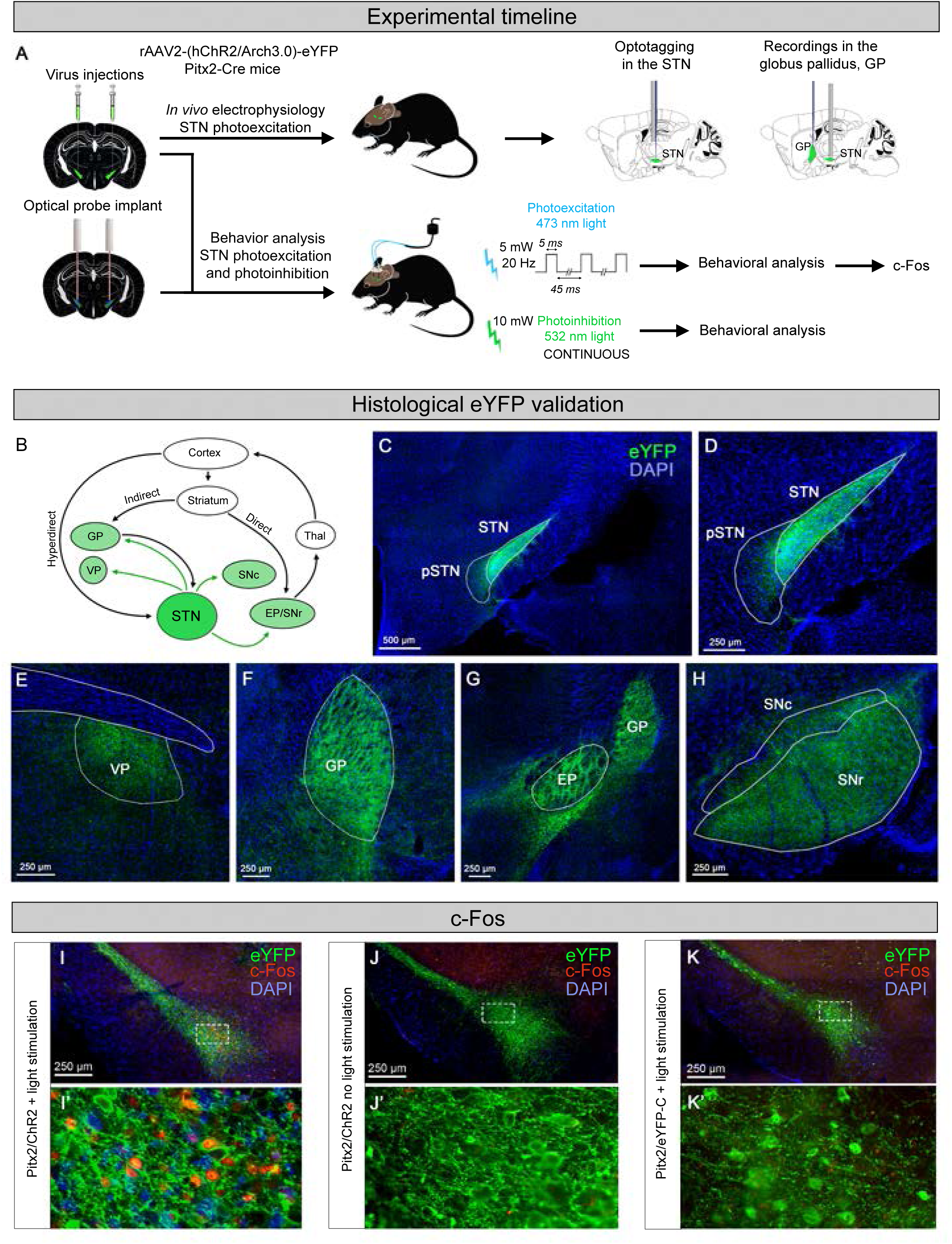
Expression of optogenetic constructs in the STN of Pitx2-Cre mice with c-Fos induced upon photoexcitation confirms experimental setup. (A) Timeline of the experimental setup. Left: graphical representation of stereotaxic bilateral injection of rAAV carrying either a Cre-dependent Channelrhodopsin 2 or Archeorhodopsin 3.0 fused with eYFP into the STN of Pitx2-Cre mice; Right: overview of the electrophysiological and behavioral experiments. (B) Simplified representation of the basal ganglia circuitry and associated structures. (C-D) Representative image of eYFP fluorescence (green) in STN cell bodies in Pitx2-Cre mice, including a close-up (D); nuclear marker (DAPI, blue); scale, 500 µm (C) and 250 µm (D). (E-H) Representative images of coronal sections showing eYFP fluorescence (green) in STN neurons target areas projections: (E) VP, (F) GP, (G) GP and EP and (H) SNr and SNc; nuclear marker (DAPI, blue); scale, 250 µm. (I) Example image showing eYFP (green) and c-Fos expression in the STN of Pitx2/ChR2 mice upon 120s photostimulation *in vivo*. (I’) High magnification showing colocalization of eYFP and c-Fos positive neurons. (J, J’, K, K’). None of the two controls, Pitx2/ChR2 without light stimulation and Pitx2/eYFP-C with light stimulation, showed c-Fos expression in the STN. For (H), (I) and (J), the white rectangles represent the corresponding high magnification pictures. Abbreviations: STN, subthalamic nucleus; pSTN, para-subthalamic nucleus; VP, ventral pallidum; GP, globus pallidus; EP, entopeduncular nucleus; SNr, substantia nigra *pars reticulata*; SNc, substantia nigra *pars compacta*; Thal, thalamus.

Different cohorts of mice were used for electrophysiological recordings and behavioral analyses, respectively (experimental outline, Figure 1A; statistical analysis, Supplementary Table 1). Prior to sacrifice, behavioral Pitx2/ChR2 and Pitx2/eYFP-C mice underwent prolonged photostimulation and brains were dissected for histological validation of c-Fos, an indicator of induced neuronal activity. Upon completed functional analyses, all mice were analyzed histologically. Reporter expression, injection site and cannula placement were validated. Mice that displayed strong cellular eYFP labeling throughout the extent of the STN and optical cannulas placed immediately above the STN were included in the statistical analyses.

Histological analysis of eYFP fluorescence in the STN and projection target areas (Figure 1B) verified the selectivity of eYFP labeling in the STN structure over surrounding structures (Figure 1C-D). The entire extent of the STN was labeled, and within STN neurons, eYFP was detected throughout the cell body. No eYFP was detected in adjacently located structures, apart from weakly labeled fibers in the para-STN. Strong YFP labeling was identified in projections reaching the ventral pallidum, entopeduncular nucleus (EP, corresponding to GPi in primates), GP (GPe in primates), SNr and substantia nigra *pars compacta* (SNc) (Figure 1E-H). Further, Pitx2/ChR2 mice that had received photostimulation showed robust c-Fos labeling in the STN (Figure 1I-I’) while no c-Fos labeling was detected in mice not stimulated (Figure 1J-J’) or in Pitx2/eYFP-C control, even upon photostimulation (Figure 1K-K’). Thus, the STN and its target areas express the optogenetic constructs and allow neuronal activation upon photostimulation.

### Optogenetic activation excites STN neurons and induces post-synaptic responses

To ensure firing and assess connectivity upon optogenetic activation of the STN, Pitx2/ChR2 mice were analyzed in two electrophysiological paradigms. Light-evoked responses of both the STN itself and one of its main target areas in motor control, the GP, were studied by *in vivo* single cell electrophysiological recordings upon optogenetic stimulation of the STN. First, an optotagging protocol (Figure 2A) was used to stimulate and record within the STN. To observe the reaction of STN neurons to photostimulation, peri-stimulus time histograms (<u>PSTH protocol</u>, 0.5 Hz, 5 ms bin width, 5-8 mW) were created by applying a 0.5 Hz stimulation protocol for at least 100 seconds. Action potentials in ChR2-positive STN cells were successfully evoked by STN photostimulation (Figure 2B).

**Figure 2.**
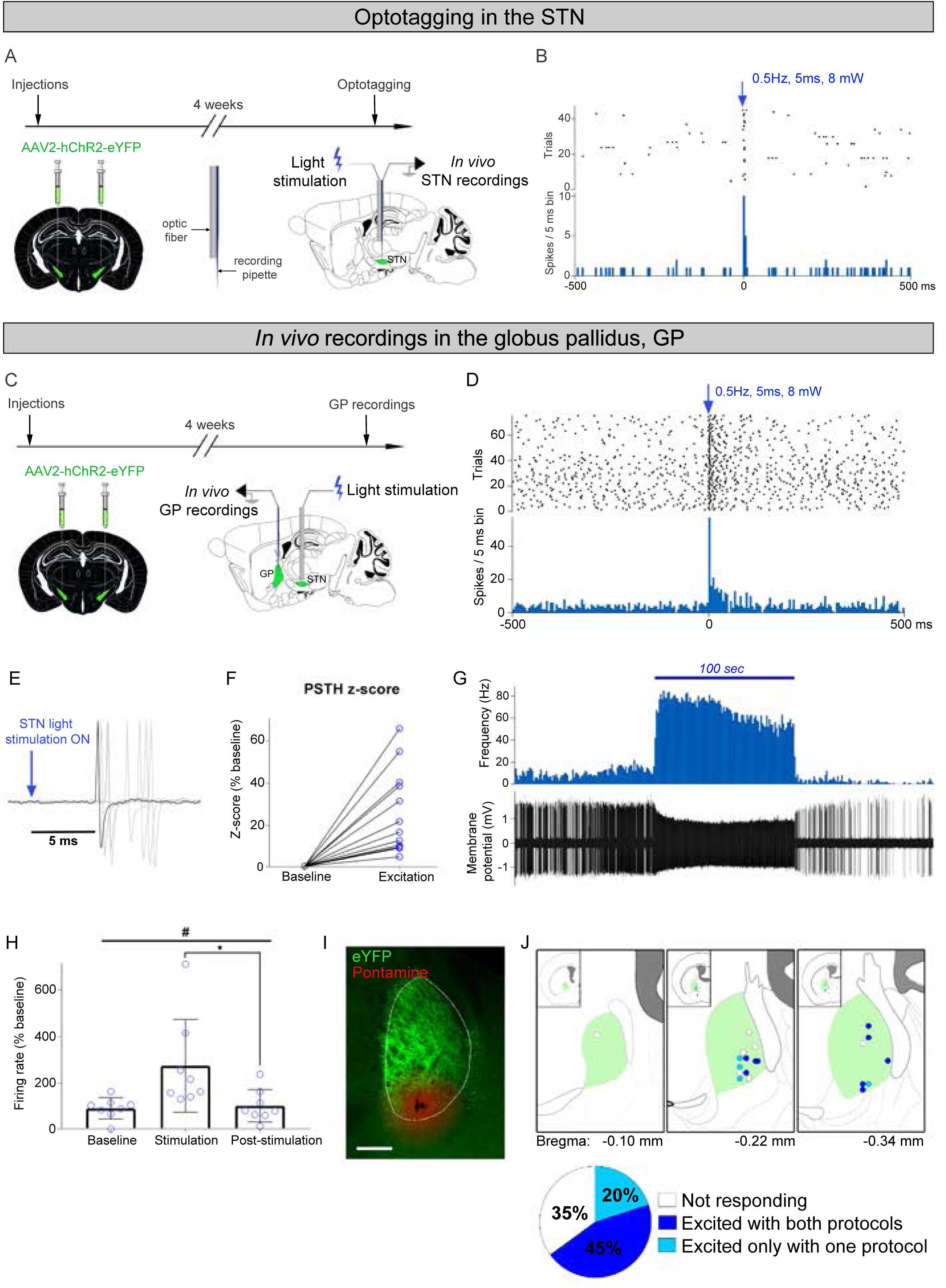
Optogenetic activation excites STN neurons and induces post-synaptic responses within the basal ganglia. (A) Procedure for STN optotagging experiments. (B) Example of a PSTH and raster (trials) of a neuron excited upon light stimulation. (C) Procedure of GP recordings experiments with STN photostimulation. (D) Example of a raster and PSTH of a GP neuron excited by STN photostimulation. (E) Overlapped recorded spikes of a GP neuron triggered by STN photostimulation during the PSTH protocol. (F) z-score of 12 GP neurons excited during the PSTH protocol. (G) Example of an excited GP neuron during the Behavioral protocol with the frequency and the recording trace. (H) Firing rate of excited GP neurons before, during and after 100 seconds of light stimulation as shown in g. Results showed a significant difference in the firing rate between “Stimulation” and “Post-stimulation”, #p<0.05, *p<0.05. (I) eYFP fibers in the GP and sky blue pontamine deposit (red dot) at the last recorded coordinate (scale, 200 µm). (J) Reconstructed mapping of GP recorded neurons at three different antero-posterior levels (N=20 neurons recorded in 3 mice). Dark blue dots indicate neurons excited during the two different protocols (PSTH and Behavioral protocols), light blue dots indicate neurons excited during one of the two protocols and white dots correspond to neurons which did not respond to the light. The pie chart shows that about two third of GP neurons were excited upon STN photostimulation. Blue arrow in B, D and E shows STN photostimulation. Abbreviations: STN, subthalamic nucleus; GP, globus pallidus.

Next, recordings were performed within the GP structure upon two different stimulation paradigms applied in the STN (Figure 2C). Firstly, the PSTH protocol was implemented which evoked action potentials in the majority of the recorded GP neurons (Figure 2D-F). This finding verified functional connectivity between the STN and GP. Next, based on previous confirmation of glutamate release upon optogenetic excitation (Viereckel et al., 2018), a photostimulation protocol designed to drive STN excitation in freely-moving animals was tested (<u>Behavioral protocol</u>, 20 Hz, 5 ms pulses, 5-8 mW; 100 seconds). Recordings in the GP identified increased frequency and firing rate of GP neurons for the whole duration of the stimulation, after which they returned to normal (Figure 2G-H). This finding validated post-synaptic activity in the GP upon application of the protocol intended for use in upcoming behavior analyses.

Pontamine sky blue staining confirmed the positioning of the recording electrodes within the GP structure and showed that all responding GP neurons were distributed in the central/medial aspect of the GP structure (Figure 2I-J). Quantification showed that 45% of the recorded neurons responded with excitation to both the PSTH protocol and the Behavioral protocol while 20% responded to only one of the protocols and the remaining 35% did not respond at all (Figure 2J). Due to the low spontaneous activity of STN neurons, similar recordings were not performed to verify the inhibitory effect of Arch3.0. Having confirmed optogenetics-driven excitation of STN neurons and their post-synaptic responses, behavioral motor effects upon optogenetic STN activation and inhibition were next assessed.

### Optogenetic excitation and inhibition of the STN induce opposite effects on both horizontal and vertical locomotion

First, Pitx2/ChR2 and Pitx2/Arch mice and their respective controls were analyzed in the open field test, allowing a qualitative and quantitative measurement of motor activity. The open field test also allows analysis of additional functions such as exploration, fear and anxiety. Mice were allowed to explore the apparatus for 5 minutes before starting the experimental protocol. This habituation period serves to avoid effects due to separation stress and agoraphobia which can induce thigmotaxis. Following habituation, mice were tested in a single 20-minute session composed of four alternating photostimulation-off (OFF) and photostimulation-on (ON) epochs (OFF-ON-OFF-ON) using laser sources providing blue (for Pitx2/ChR2 mice and controls) and green (for Pitx2/Arch mice and controls) light.

During OFF epochs, primarily horizontal and vertical movements were observed, with short periods of grooming. To address motor effects upon STN excitation, the photoexcitation paradigm called Behavioral protocol was applied. In response to STN photostimulation (ON), Pitx2/ChR2 mice responded with a significant reduction in vertical activity, known as rearing (Figure 3A). Also, horizontal activity, measured as distance moved, speed and time spent moving was decreased upon STN-photostimulation (Figure 3B-D). By decreasing these motor parameters upon optogenetic activation of the STN (ON compared to OFF), the expected role of STN excitation in suppressing locomotion could be verified experimentally.

**Figure 3.**
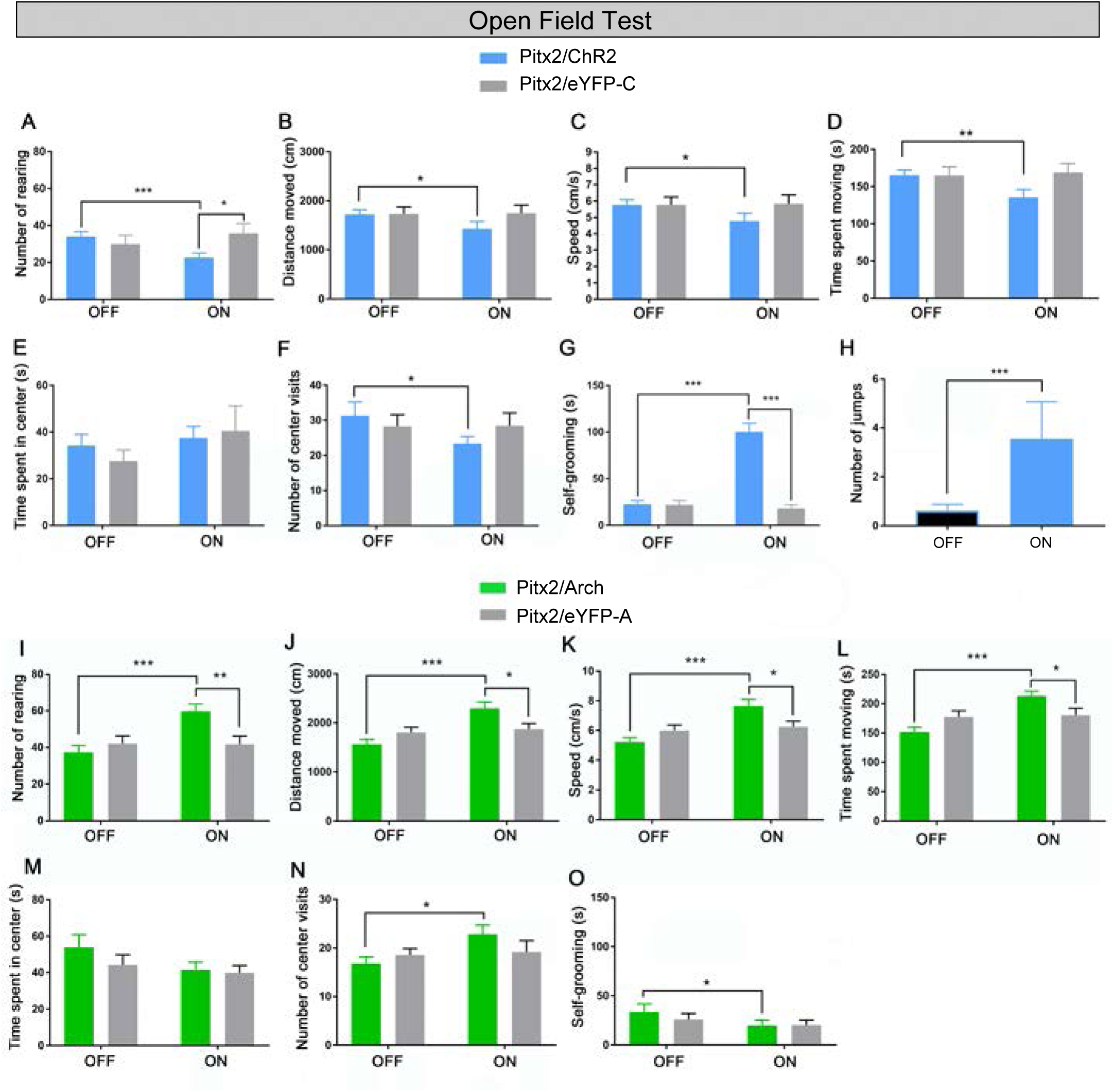
Optogenetic activation and inhibition of the STN induce opposite effects on both horizontal and vertical locomotion. (A-E) In the open field test, several behaviors representative of vertical and horizontal locomotion were recorded in Pitx2/ChR2 mice and their corresponding Pitx2/eYFP-C control mice: rearing (A), distance moved (B), speed (C), time spent moving (D), times spent exploring the center (E) and center visits (F). (G-H) Graphs showing the time spent grooming (G) and the number of jumps (H). (I-N) The same behavioral parameters were recorded in Pitx2/Arch and Pitx2/eYFP-A control mice: rearing (I), distance moved (J), speed (K), time spent moving (L), time spent exploring the center (M) and visits to the center (N). Time spent grooming (O) was also recorded. All data are presented as average for the two ON and OFF epochs ± SEM, *p<0.05, **p<0.01, ***p<0.001.

The time spent exploring the center of the arena, a measure of anxiety, was not affected by the stimulation (Figure 3E), but the number of exploratory visits to the center was significantly lower during ON epochs (Figure 3F). In addition, some Pitx2/ChR2 mice displayed abnormal gait upon photoexcitation, characterized by sliding or slipping on the floor or backwards movements as indication of a lack of balance, while some showed a prominent jumping behavior (Figure 3H).

Contrary to STN activation in Pitx2/ChR2 mice, but in accordance with the anticipated role of the STN in motor control, continuous optogenetic STN inhibition in Pitx2/Arch mice resulted in increased horizontal and vertical movement (Figure 3I-L). In contrast to STN excitation in Pitx2/ChR2 mice, while not affecting the time spent exploring the center (Figure 3M), STN inhibition increased exploratory activity, visible by a higher number of visits to the center during ON epochs (Figure 3N).

Further pinpointing the opposite effects on motor parameters by STN excitation *vs* inhibition, also grooming behavior was altered in opposite directions by the optogenetic manipulations. Excitation of the STN induced significant face-grooming behavior in the Pitx2/ChR2 mice (Figure 3G, Video clip 1). In contrast, the naturally occurring grooming behavior was lower in Pitx2/Arch mice upon photoinhibition than during OFF epochs (Figure 3O). In rodents, grooming is an innate behavior which follows a distinct pattern, referred to as the cephalo-caudal rule, covering the extent of the body but starting in the face region (Berridge et al., 2005; Fentress, 1988). The grooming displayed by Pitx2/ChR2-eYFP mice was directly associated with STN-photostimulation. While grooming was not continuous throughout each stimulation phase in all mice, it was initiated shortly upon STN-photostimulation to most often be observed in numerous episodes of separate activity. Further, the observed behavior did not follow the cephalo-caudal grooming rule, but was displayed as repetitive strikes by both front paws tightly in the face area, primarily around the nose (Video clip 1).

Together, these results confirm the importance of the activational level of the STN in horizontal and vertical movement. The data also identify a direct correlation between STN excitation and stereotyped grooming.

### Unilateral excitation and inhibition of the STN induce opposite rotations

To further assess the impact of optogenetic manipulations on motor activity, unilateral STN photostimulation was performed in Pitx2/ChR2 and Pitx2/Arch mice (Figure 4A). Since unilateral activation or inhibition of the STN is expected to give rise to rotational rather than forward movement, this was scored by comparing ipsilateral and contralateral rotations. Indeed, STN-photostimulation induced a strong rotational behavior. Pitx2/ChR2 mice made a significantly higher number of ipsilateral than contralateral rotations during ON epochs (Figure 4B-C). In contrast, Pitx2/Arch mice made significantly more contralateral than ipsilateral rotations during ON epochs (Figure 4E-F). No rotational behavior was detected for control mice during either ON or OFF epochs (Figure 4D and 4G). The results confirm the importance of the STN for coordinated motor output by demonstrating that manipulation of one of the two STN structures is sufficient to produce measurable rotation effects. No abnormal gait, jumping or grooming was observed during unilateral stimulations.

**Figure 4.**
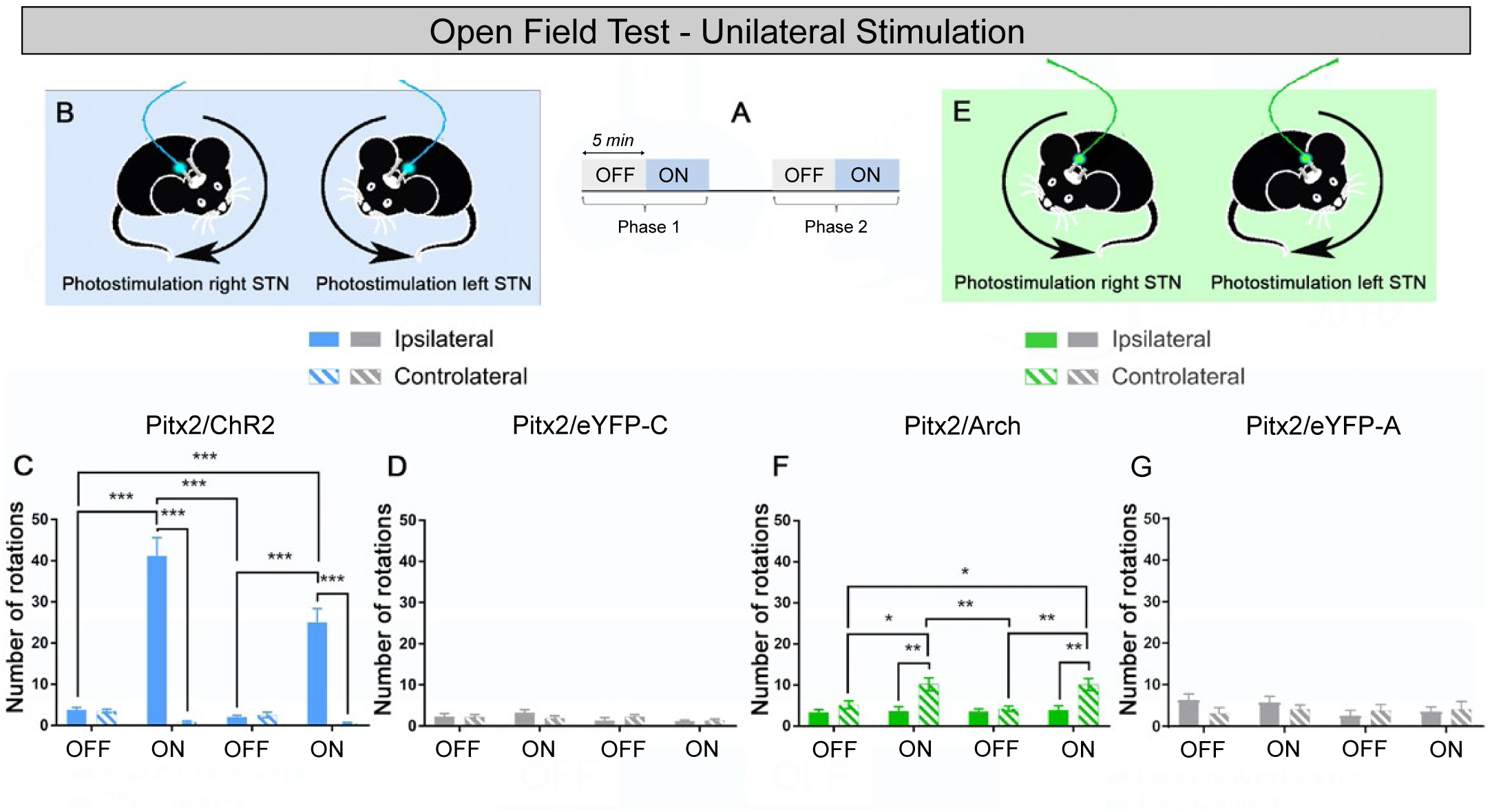
Unilateral inhibition and excitation of the STN induce opposite rotations. (A) Schematic representation of the open field test with unilateral stimulation. (B) Graphic representation of types of body rotations displayed by Pitx2/ChR2 mice depending on stimulated hemisphere. (C-D) Number of ipsilateral and contralateral rotations in Pitx2/ChR2 mice (C) and Pitx2/eYFP-C control mice (D) upon STN photoexcitation. (E) Graphic representation of types of body rotations displayed by Pitx2/Arch mice depending on stimulated hemisphere. (F-G) Number of ipsilateral and contralateral rotations in Pitx2/Arch mice (F) and Pitx2/eYFP-A control mice (G) upon STN photoinhibition.

### Optogenetic STN-activation disrupts motor coordination in the rotarod test

The strong reduction in locomotion together with gait abnormality and prominent grooming behavior observed during the open field test of Pitx2/ChR2 mice, but not Pitx2/Arch mice, prompted further analysis of the impact of STN excitation on motor coordination. Pitx2/ChR2 and control mice were next analyzed in both the rotarod and the beam walk tests. First, the ability to sustain on-going motor coordination was assessed in the rotating rotarod. Upon training to master maintained walking on the rotating rod throughout a two-minute session, mice were exposed to STN photostimulation in four OFF/ON trials (Figure 5A-B). Pitx2/ChR2 and control mice all managed to stay on the rod throughout the entire length of the OFF trials. During ON trials, photostimulation was applied 5 seconds after the mouse was properly walking on the rod. All Pitx2/ChR2 mice fell off the rod within seconds after STN excitation was initiated, giving rise to a strikingly short latency to fall (Figure 5C, Video clip 2). In contrast, all control mice maintained their rod-walking undisturbed (Figure 5D). The disrupting effect on motor coordination in Pitx2/ChR2 mice was transient and displayed specifically upon photoexcitation, with completely restored coordination during subsequent OFF trial (Figure 5C). Further, since bilateral excitation in the open field test was associated with grooming, specific attention was directed to observe if this behavior was manifested while walking on the rotating rod. However, it was clear no grooming behavior was displayed (Video clip 2). Thus, the loss of motor coordination on the rotating rod was independent of grooming.

**Figure 5.**
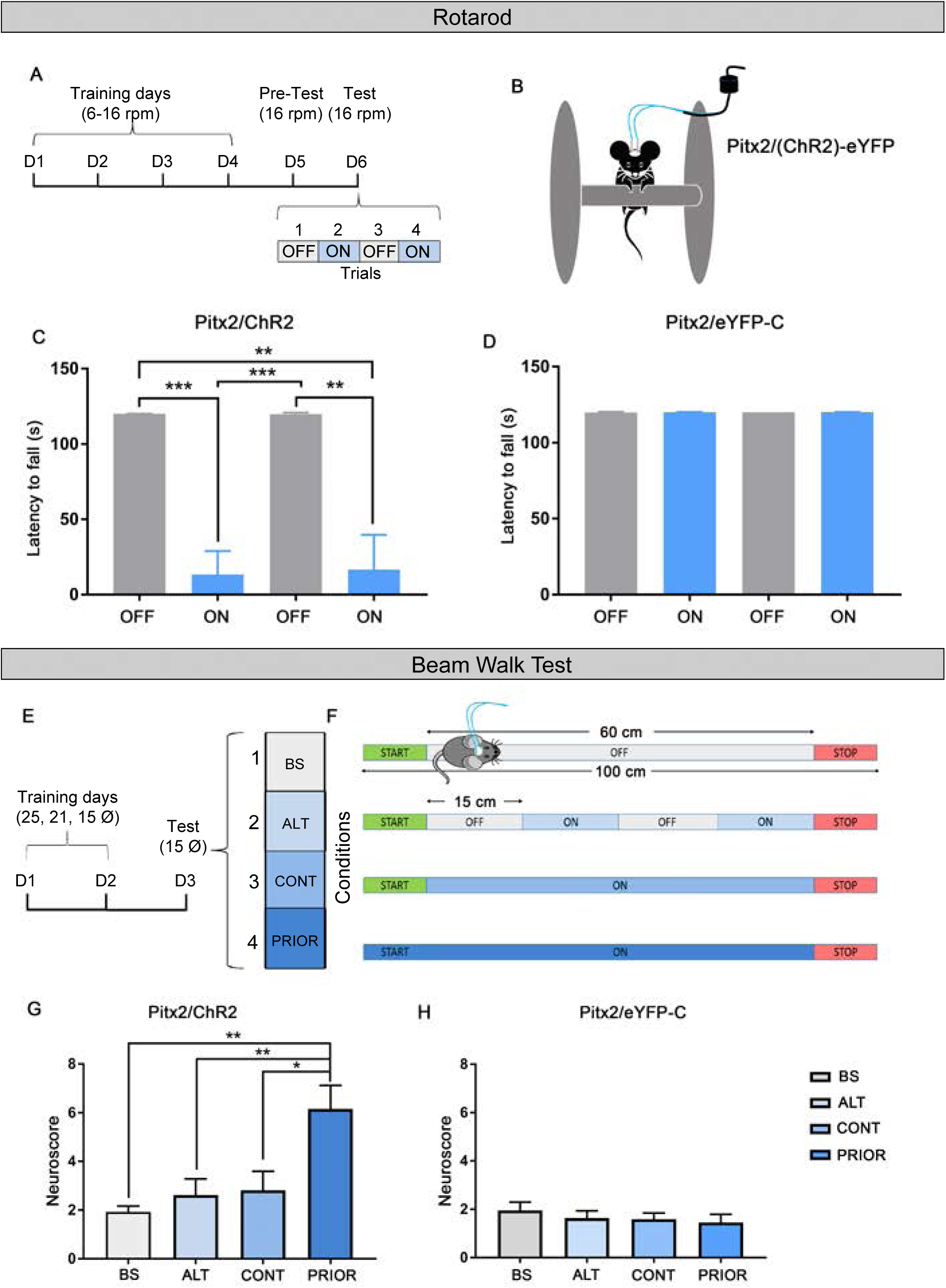
Optogenetic STN-activation disrupts motor coordination. (A) Schematic representation of the fixed speed rotarod assay timeline. (B) Graphical representation of the rotarod assay, with the mouse receiving bilateral optical stimulation during ON trials. (C) Latency to fall from the rotating rod for Pitx2/ChR2 is expressed as mean ±SEM, **p<0.01, ***p<0.001 and (D) for Pitx2/eYFP-C control mice, ±SEM, p=0.1989. (E) Schematic representation of the beam walk test time line. (F) Schematic representation of the tested conditions: “*Baseline*” (BS), “*Alternated Light*” (ALT), “*Continuous Light*” (CONT) and “*Prior Stimulation*” (PRIOR). (G-H) The neuroscore is calculated from a scoring scale (see Supplementary Table 2) and is expressed as mean ±SEM, *p<0.05, **p<0.01.

The potent but transient effect on motor coordination upon STN excitation was subsequently assessed in the beam walk test (Figure 5E-F). Upon training, three conditions were tested in order to determine how the excitation of the STN would influence the motor coordination necessary to cross the beam: alternated OFF and ON epochs (referred as ALT), continuous stimulation along the beam (referred as CONT) or light stimulation starting few seconds before and during the cross of the beam (referred as PRIOR) (Figure 5E). The results were analyzed and summarized in a neuroscore pre-defined rating scale (Supplementary table 2). The maximum score was 10 points for each condition (5 points for latency to cross + 5 points for hind-foot slips). The lower is the score, the better is the motor performance. During the last condition only (PRIOR), motor coordination was altered to significantly increase the number of paw slips and latency to cross (Figure 5G). This finding suggests that a prolonged STN excitation is necessary to impair motor coordination in the beam walk test. Further, no grooming or jumping behavior was observed. No effect upon photostimulation was observed in control mice (Figure 5H). Taken together, the rotarod and beam walk tests identify a direct association between STN excitation and impairment of motor coordination when mice are engaged in performing challenging motor tasks.

## Discussion

The role of the STN in normal motor regulation has long been assumed as established. However, experimental evidence is sparse. Since the execution of intended movement depends on several different parameters to be fully functional, a solid identification of how each of these are regulated should contribute to decoding the complexity of motor function. Forward locomotion, gait, balance and motor coordination are all of critical importance to achieve intended movement, and their disturbance manifested in motor disorders, including PD, supranuclear palsy, Huntington’s disease and hemiballismus, all in which STN dysfunction is implicated. In this study, we found that optogenetic STN excitation was generally correlated with significant reduction in locomotor activity, while in contrast, optogenetic STN inhibition enhanced locomotion, just as the classical model of the basal ganglia would predict. Optogenetic excitation *vs* inhibition thus verify the basal ganglia model in terms of locomotion. However, we also found that these correlations were not true for all types of movement. Upon optogenetic excitation of the STN in the open field test, jumping and self-grooming behaviors were induced, not reduced. Curiously, this was only observed in a non-challenging environment, not when mice where engaged in advanced motor tasks. Together, these findings indicate that detailed analysis of the behavioral roles driven by the STN is likely to enhance knowledge of motor control and its disease, beyond what might be expected based on classical models.

We recently showed that *in vivo* optogenetic stimulation of the STN at 20 Hz, which is within the range of stimulation frequency that excites rodent STN neurons during normal conditions, in Pitx2/ChR2 mice was sufficient to release measurable amounts of glutamate in the GP (Viereckel et al., 2018). Based on this finding, the Behavioral protocol for photoexcitation of the STN used throughout the behavior analyses of Pitx2/ChR2 mice was implemented at this frequency. This photoexcitation protocol was also validated by *in vivo* electrophysiological recordings performed in intact anaesthetized mice. Further, to ensure firing and assess connectivity upon optogenetic activation of the STN, Pitx2/ChR2 mice were also analyzed for optogenetically evoked responses using peri-stimulus time histograms by applying a 0.5 Hz stimulation protocol. The results firmly demonstrated that action potentials were reliably evoked in both the STN and the GP, a critical STN target area. Electrophysiological recordings showed that GP neurons were excited already upon a 0.5 Hz photostimulation of the STN structure, but also that the 20 Hz Behavioral protocol gave rise to the majority of excitatory responses. With these validations, the behavioral assessments using ChR2 to excite the STN neurons in their natural brain network rest upon electrophysiological confirmation within both the STN itself and GP. However, additional recordings would be required to fully pinpoint the range of target areas that mediate the different behavioral responses observed, as additional brain structures beyond the GP might be involved. Verifying the expected innervation pattern, STN-derived eYFP-positive projections were identified in the GP, EP, SNr and SNc, structures that may all be involved in the observed behaviors. Another limitation of the study is the lack of electrophysiological recordings upon STN inhibition in Pitx2/Arch mice, which were not performed in the current setup due to the low spontaneous activity of the STN *in vivo*.

When addressing behavior, we initially used both optogenetic STN excitation (ChR2) and inhibition (Arch) to enable the comparison of results. This distinct experimental separation into STN excitation *vs* STN inhibition clearly caused different effects on several aspects of motor function with reduced forward locomotion and rearing upon STN excitation, and enhancement of both parameters upon inhibition. These findings firmly verify the opposing roles executed upon STN excitation and inhibition, with a mirror effect displayed by these opposite types of stimulations. Beyond the comparison with STN inhibition, the analyses also identified several unexpected behaviors upon STN excitation that deserved additional attention. In contrast to the strong history of STN inactivation by surgical lesioning, optogenetics offers the novelty of controlled excitation. This is an important experimental advantage because in its natural network, STN is excited by cortical projections via the hyperdirect pathway and is more or less inhibited via the indirect pathway, and in turn, exerts excitatory influence over its targets. Thus, while STN inactivation and inhibition are both of major focus in PD-related research in order to find ways to override pathological STN hyperactivity and promote movement, the naturally occurring regulatory role of the STN upon excitation has been far less explored. Here, we could observe that optogenetic excitation of the STN in Pitx2/ChR2 mice caused jumping and self-grooming behavior in the open field test, and when explored further in more advanced motor tests, Pitx2/ChR2 mice displayed loss of balance and motor coordination. These behavioral displays were observed immediately upon STN excitation in Pitx2/ChR2 mice. Notably, the mice did not lose balance or coordination due to induced jumping or self-grooming, as these types of behavior were not detected when mice were performing in the more advanced tests, the rotating rotarod and the beam walk. Thus, it seems that when mice are engaged in challenging motor tasks, STN excitation causes interruption leading to loss of coordination, while when they are freely exploring, STN excitation causes grooming, jumping and/or loss of normal gait. Further, the strong impact of STN excitation on balance and coordination likely reflects a disturbed ability to maintain position, due to reduced movement, in challenging conditions where flexible motor output is required. One recent study identified that on-going licking behavior in mice was interrupted by optogenetic excitation of STN neurons (Fife et al., 2017). While licking behavior is distinct from the type of motor coordination required to master the rotating rotarod, it might be noteworthy that behaviors that ought to require substantial motivation or engagement are disturbed when the STN is excited.

Previous reports have shown that inactivation of STN function, either by surgical STN lesioning (Centonze et al., 2005; Heywood & Gill, 1997; Parkin et al., 2001; Rizelio et al., 2010) or by conditional knockout of the *Vglut2* gene in the STN of Pitx2-Cre mice (Schweizer et al., 2014) significantly elevated the level of locomotion. These findings, which provided support to the basal ganglia model in which STN inhibition is postulated to increase movement, are in accordance with the current study in which we identify that optogenetic inhibition of the STN causes hyperkinesia while optogenetic excitation causes hypokinesia during non-pathological conditions. The opposite effects of STN excitation and inhibition observed here also lend indirect support for the inhibition hypothesis of DBS, albeit during non-parkinsonian conditions. Despite the current application of STN-DBS as treatment of PD, the mechanisms underlying its symptom-alleviating effects are still not entirely understood. It is currently debated whether the neural elements are excited or inhibited by the stimulations (Chiken & Nambu, 2016; Deniau et al., 2010). One model of DBS mechanism involves the depolarization of axons and inhibition of cell bodies (Florence et al., 2016; Vitek, 2002), while another model proposes excitation of afferent, efferent and passing fibers (Kringelbach et al., 2007) with increased firing (Hashimoto et al., 2003) and glutamate release in basal ganglia output areas (Windels et al., 2000). Studies based on optogenetics in parkinsonian animal models have led support to both the inhibition and excitation hypothesis of STN-DBS (Gradinaru et al., 2009; Sanders & Jaeger, 2016; Yoon et al., 2014). Recently, a study using the ultrafast opsin CHRONOS, aiming at creating an animal model of optogenetic STN-DBS, showed that parkinsonian movement disability was improved to a similar degree as when electrical STN-DBS was applied, further strengthening the correlation between the STN and motor output in the parkinsonian model (Yu et al., 2020). In PD and DBS contexts, our results showing opposite locomotor effects upon optogenetic STN excitation and inhibition, with enhanced locomotion upon STN inhibition, thus provide indirect support for the inhibition hypothesis of STN-DBS. However, more evidently, the current results clearly identify effects beyond locomotion, as gait, balance and motor coordination parameters are all affected by STN manipulations. Future studies might be of interest in which a parkinsonian model such as 6-OHDA toxicity is applied in the Pitx2/ChR2 and Pitx2/Arch mice in order to unravel the effects upon optogenetic excitatory *vs* inhibitory stimulations on pathological motor performance.

*Pitx2* gene expression is a characteristic property of STN neurons, with expression starting early in STN development and persisting throughout life (Dumas & Wallén-Mackenzie, 2019; Martin et al., 2004; Schweizer et al., 2016; Skidmore et al., 2008). In previous work, we could verify the selectivity of Cre recombinase expressed under control of *Pitx2* regulatory elements (Pitx2-Cre) in driving double-floxed ChR2 constructs to the STN, with photostimulation-induced post-synaptic currents and glutamate release observed in STN target areas of the basal ganglia (Schweizer et al., 2014; Viereckel et al., 2018), as well as its selectivity for targeting the floxed *Vglut2* allele specifically in the STN (Schweizer et al., 2014, 2016). Thus, the transgenic mouse line used here has been well validated for its selectivity in driving recombination of floxed alleles in the STN. As discussed above, one main question in the pre-clinical and clinical STN fields, not least in the context of STN-DBS, has been the contribution of the STN relative afferent and efferent projections in mediating the behavioral roles assigned the STN (Gradinaru et al., 2009; Hamani, 2004; Sanders & Jaeger, 2016; Tian et al., 2018). Further, the STN structure itself has remained elusive, with regional distribution proposed according to a tripartite model which subdivides the primate STN into three major domains, of which the dorsal-most domain of the STN is postulated to mediate motor functions (Haynes & Haber, 2013; Lambert et al., 2012; Yelnik et al., 2007). The *Pitx2* gene is expressed throughout the entire STN structure (Dumas & Wallén-Mackenzie, 2019; Wallén-Mackenzie et al., 2020), and with the present study, we have shown that optogenetic excitation and inhibition of the STN clearly drive distinct motor effects. In a parallel study, we have recently addressed the anatomical organization of the mouse STN using single cell transcriptomics followed through with multi-fluorescent histological analysis. This combined analysis allowed us to subdivide the mouse STN into three main domains based on distinct markers, gene expression patterns (Wallén-Mackenzie et al., 2020). With the current optogenetics-based data at hand revealing regulatory roles of the STN in several motor parameters, including locomotion, balance, self-grooming and motor coordination, future studies exploring the possibility that regional distribution of neurons within the STN structure might be differentially involved in mediating these functions should be of interest. While the Pitx2-Cre transgene successfully directs floxing to the STN, new possibilities for spatial selectivity within the STN itself might be possible using the new markers identified (Wallén-Mackenzie et al., 2020).

For example, the strong and immediate self-grooming behavior upon optogenetic STN excitation might deserve special attention. Curiously, this unexpected behavior was only observed in a non-challenging environment, not when mice where engaged in complex motor tasks. While self-grooming is a measurable display of motor output, it might represent emotional or associative manifestations (Smolinsky et al., 2009; Wahl et al., 2008). Beyond PD, the STN is a new target for DBS treatments in cognitive and affective disorders, including obsessive compulsive disorder (OCD) (Mallet et al., 2008). Stereotypy, such as repetitive and excessive grooming, is a behavioral criteria for rodent models of OCD, and is commonly interpreted as a rodent equivalent to compulsivity in patients with OCD (Kalueff et al., 2016). Indeed, high-frequency stimulation of the STN has been demonstrated as effective in the treatment compulsive-like behaviors in rodents (Klavir et al., 2009; Winter et al., 2008) and non-human primates (Baup et al., 2008). In this context, our results on repetitive grooming activity upon optogenetic STN excitation suggest that the positive effects of STN-DBS in treatment of compulsivity in OCD might be due to an inhibitory effect on STN neurons. Such an effect could result in a modulatory control of the basal ganglia output areas. In accordance with this hypothesis, lesions of the GP impaired grooming (Cromwell & Berridge, 1996), while pharmacological inactivation of the EP and GP exerted an anti-compulsive effect on quinpirole-sensitized rats (Djodari-Irani et al., 2011). On the other hand, optogenetic activation of the direct pathway induced repetitive behaviors (Bouchekioua et al., 2018), while repeated optogenetic stimulation of the cortico-striatal pathway triggered OCD-like self-grooming (Ahmari et al., 2013). In our study, photoexcitation-evoked grooming activity in Pitx2/ChR2 mice did not follow the cephalo-caudal rule but was restricted to the face area, and the grooming ceased with the removal of the stimulation. This result suggests a difference in the control of grooming between STN and striatum, which may act in parallel to integrate signals to the basal ganglia output areas to control abnormal actions such as repetitive behaviors. The neurological underpinnings of the STN in compulsive behavior will need further attention. However, it is evident from the present results that STN excitation drives self-grooming of the face area. With new markers that define domains within the mouse STN (Wallén-Mackenzie et al., 2020), optogenetic dissection of the putative role of each distinct domain in different aspects of motor function, including grooming, should be of interest to pursue.

## Conclusion

Locomotion, balance and coordination are critical motor features required to enable and sustain voluntary movement. The experimental approach used in this study allowed the firm identification of a clear correlation between the STN and regulation of each of these pivotal motor functions. The results presented provide experimental evidence supporting the predicted role of the STN according to classical basal ganglia models, and identify an unexpected role of the STN in self-grooming behavior. Further studies will be required to pinpoint mechanistic underpinnings and circuitry components involved in mediating each of these distinct behaviors.

**Video clip 1**

Video clip of the open filed test showing first a Pitx2/ChR2 mouse in the arena without light stimulation, and second, the same mouse grooming seconds after STN photoexcitation initiated. A close-up video shows the grooming behavior from the side.

**Video clip 2**

Video clip of the rotarod test showing first a Pitx2/ChR2 mouse walking on the rod without STN photoexcitation (OFF trial) and second, the same mouse falling off the rod few seconds after STN photoexcitation initiated (ON trial, repeated one time at the end of the video clip).

## Supplementary tables

**Supplementary Table 1**

Detailed description of the statistics used for the different experiments with reference to each figure where the data is displayed.

Abbreviations: STN, subthalamic nucleus; GP, globus pallidus; OFT, open field test; BWT, beam walk test.

**Supplementary Table 2**

Table shows the parameters used to calculate the neuroscore for the beam walking test. Based on Korenova et al., 2009 and adapted for mice (Korenova et al., 2009).

## Acknowledgements

We thank Professor James Martin, Baylor College of Medicine, Houston, Texas, USA, for generously providing the Pitx2-Cre transgenic mouse line. Members of the Mackenzie Lab are thanked for constructive feedback throughout this study.

## Author contributions

A.G: Investigation, Formal analysis, Visualization, Writing – original draft; G.P.S: Investigation, Formal analysis, Methodology, Visualization, Writing – review and editing; F.G: Supervision; Å.W.M: Conceptualization, Funding acquisition; Project administration, Supervision; Writing - original draft, review and editing.

